# MIST: an interpretable and flexible deep learning framework for single-T cell transcriptome and receptor analysis

**DOI:** 10.1101/2024.07.05.602192

**Authors:** Wenpu Lai, Yangqiu Li, Oscar Junhong Luo

## Abstract

Joint analysis of transcriptomic and T cell receptor (TCR) features at single-cell resolution provides a powerful approach for in-depth T cell immune function research. Here, we introduce a deep learning framework for single-T cell transcriptome and receptor analysis, MIST (Multi-Insight for T cell). MIST features three latent spaces: gene expression, TCR, and a joint latent space. Through analyses of antigen-specific T cells and T cells related to lung cancer immunotherapy, we demonstrate MIST’s interpretability and flexibility. MIST easily and accurately resolves cell function and antigen-specificity by vectorizing and integrating transcriptome and TCR data of T cells. In addition, using MIST, we identified the heterogeneity of CXCL13^+^ subsets in lung cancer infiltrating CD8^+^ T cells and their association with immunotherapy, providing additional insights into the functional transition of CXCL13^+^ T cells related to anti-PD-1 therapy that were not reported in the original study. MIST is available at https://github.com/aapupu/MIST.

## Introduction

T cell function is greatly influenced by its T cell receptor (TCR), which recognizes antigen and conducts signals for downstream biological processes in a TCR sequence-dependent manner (*1, 2*). With the advancement of single-cell sequencing, transcriptome and TCR delineation at single-cell resolution provides in-depth understanding of T cell function (*3, 4*). Combining single-cell RNA sequencing (scRNA-seq) and single-cell T cell receptor sequencing (scTCR-seq) has been proven an effective approach to characterizing T cells in health and disease (*3, 5*). However, most current studies treat the joint analysis of scRNA-seq and scTCR-seq data as relatively separate processes, mainly focusing on the combined analysis of gene expression (GEX) and TCR based on TCR clonotypes (*5–7*). Recent deep neural network study based on language models, such as Transformer, successfully captured the core qualities of TCRs (*8*). Therefore, developing analytical tools using deep learning to further enhance the integrated analysis of T cell biology is both critical and feasible.

Recently, several algorithms and tools have been developed for the integrative analysis of scRNA-seq and scTCR-seq data, such as CoNGA (*9*), Tessa (*10*), and scNAT (*11*). CoNGA and Tessa rely on the assumption that all T cells with the same TCR clonotype would have similar transcriptome profiles. This is not necessarily the case for T cells from different individuals, tissues, and batches, or even for T cells within complex disease microenvironment (e.g., tumor). scNAT, on the other hand, uses deep learning to merge scRNA-seq and scTCR-seq data into a joint latent space, facilitating downstream analysis by digitizing the combined features. However, scNAT only provides a joint latent representation of the multimodal data without offering separate latent representations for GEX and TCR, and the model lacks interpretability.

Here, we introduce MIST (Multi-Insight for T cell), an interpretable and flexible deep learning framework for single-T cell transcriptome and receptor analysis. Similar to scNAT, MIST’s primary goal is to merge and vectorize the features of GEX and TCR. Importantly, MIST’s modular architecture provides the latent representations for both GEX and TCR, respectively, enabling extensive and flexible downstream analysis, including clustering, batch effect removal, GEX imputation, TCR similarity calculation, and more. Additionally, MIST offers interpretability for both GEX and TCR, allowing for the exploration of specific gene expression patterns and conserved TCR motifs. MIST is available at https://github.com/aapupu/MIST as a continuously supported, open source software package.

## Results

### MIST for transcriptome and TCR integration

We developed MIST based on variational auto-encoder (VAE) architecture (*12*), which encodes input data into latent space variables following a multivariate normal distribution and decodes from these latent space variables to reconstruct the original data. MIST can compute the representation from GEX, TCR and joint latent space, respectively, for T cell functional characterization (Figure 1A left; Figure S1). MIST takes in single-T cell gene expression matrices from scRNA-seq and TCRαβ sequences from the matching scTCR-seq data as inputs, and parse them through the encoder-decoder layers with built-in self-attention mechanism (*13, 14*) for unsupervised learning (Methods). In this process, latent spaces for the transcriptome profiles (GEX) and TCR sequences are computed separately, and the joint latent space co-considering the GEX and TCR profiles is also created. Concretely, we penalize the model to learn the latent association between GEX and TCR by minimizing the distance between their representations (*15, 16*). The joint cell state is estimated as the average of both GEX and TCR latent space. Additionally, the Domain-Specific Batch Normalization (DSBN) layer of the GEX decoder helps preserve shared domain features across different GEX batches (*17, 18*). These three latent spaces thus hold the high order encoded features for GEX, TCR and the combined data, respectively, enabling multi-perspective analysis of T cells (Figure 1A right). Each set of these representations is of important utility for analyzing T cell biology with insightful interpretability. MIST is designed to process multiple single-cell transcriptome and TCR datasets simultaneously and can autonomously eliminate batch effect.

**Figure 1:**
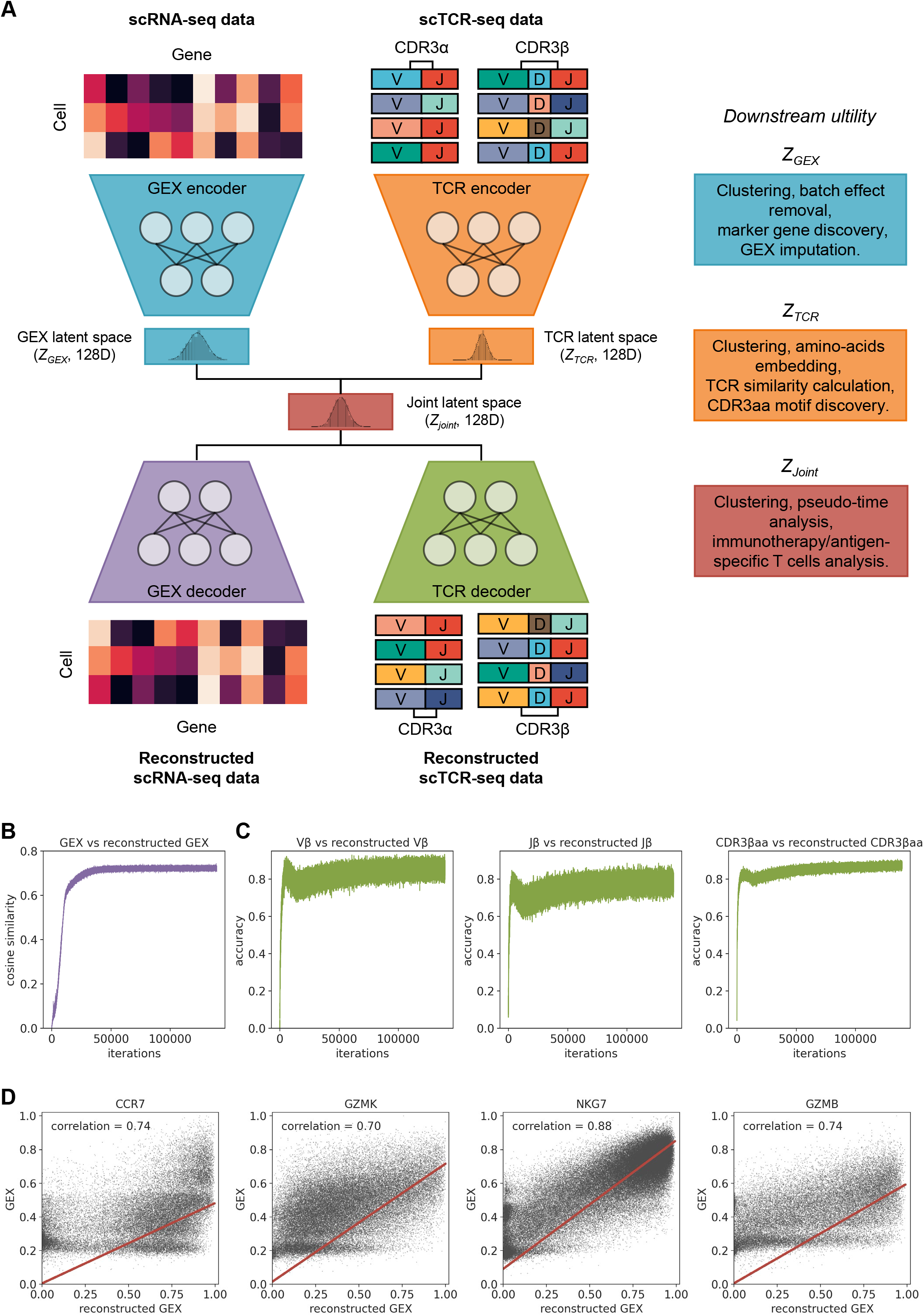
MIST model for joint scRNA-seq and scTCR-seq data analysis. A. Left: Schematics illustration of the variational auto-encoder (VAE) architecture used for merging scRNA-seq and scTCR-seq data. MIST takes scRNA-seq data (single-cell gene expression matrix, GEX) and the matching scTCR-seq data (TCRα and TCRβ sequences, TCR) as input and parse them through the encoder-decoder layers. The GEX, TCR and joint latent spaces respectively characterize GEX, TCR, and their integration, which are used for downstream functional analyses. Right: Typical usage of the GEX, TCR and joint latent spaces for respective analysis. CDR3aa: TCR CDR3 amino acid sequence. B. Line graph illustrating the increases in cosine similarity between the input GEX and the reconstructed GEX over training cycles (iterations) for the 10x_200k dataset. C. Line graphs illustrating the increases in accuracy between the input V/J gene (left and mid) and CDR3aa (right) of TCRβ and the corresponding reconstructed TCRβ segments over training cycles for the 10x_200k dataset. D. Scatterplots comparing the input and the MIST-reconstructed expression of CCR7, GZMK, NKG7 and GZMB in T cells of the 10x_200k dataset. The red lines are fitted by linear regression. The Pearson’s correlation between the input and the reconstructed expression for each gene is shown. Refer to Figure S1 and Methods for details.

### MIST facilitates intuitive single-cell GEX analysis

To evaluate whether MIST can legitimately encode transcriptome profiles and TCR sequences of T cells, we applied it to the 10x Genomics scRNA-seq and scTCR-seq dataset of approximately 200,000 CD8^+^ T cells from 4 donors (referred as the 10x_200k dataset, https://pages.10xgenomics.com/rs/446-PBO-704/images/10x_AN047_IP_A_New_Way_of_Exploring_Immunity_Digital.pdf). Our MIST model faithfully reconstructed both GEX and TCR data for T cells in the 10x_200k dataset (Figure 1B-D; Figure S2A). In more detail, we first filtered out the low-quality cells and identified the top 2,000 genes with highly variable cell-to-cell expression, as typically done in scRNA-seq data analysis workflow. Subsequently, the normalized expression counts of the top 2000 variable genes in the retained cells (as a count matrix) were fed to the GEX encoder of MIST, which features self-attention and fully connected (FC) neural network mechanism (Figure S1A, B). The unsupervised GEX VAE section of MIST aims to maximumly reconstruct the gene expression counts per cell through a series of neural network layers, in which a 128 dimensional (128D) latent space for GEX representation is computed. We then passed the 128D GEX latent representations of the 10x_200k dataset for downstream principal-component analysis (PCA), dimensional reduction visualization and cell clustering. Clearly, the encoded transcriptomic features in the GEX representations have effectively removed the scRNA-seq batch effect (inter-donor bias): cells from distinct donors assayed by different scRNA-seq experiments are well mixed (Figure 2A left). In comparison, using raw expression counts did not completely remove batch effect (*19, 20*), despite data normalization (Figure 2B-C). Eventually, by applying the Leiden clustering algorithm, the cells of the 10x_200k dataset were classified into 8 well-separated clusters (*9, 21*) (Figure 2A right). The GEX encoder can also accommodate surface protein antibody-barcode sequencing data. Projection of expression level of surface proteins critical for T cell function and subtype classification onto the Uniform Manifold Approximation and Projection (UMAP) plots showed that cell clusters derived from the GEX latent representations are functionally coherent (Figure 2D).

**Figure 2:**
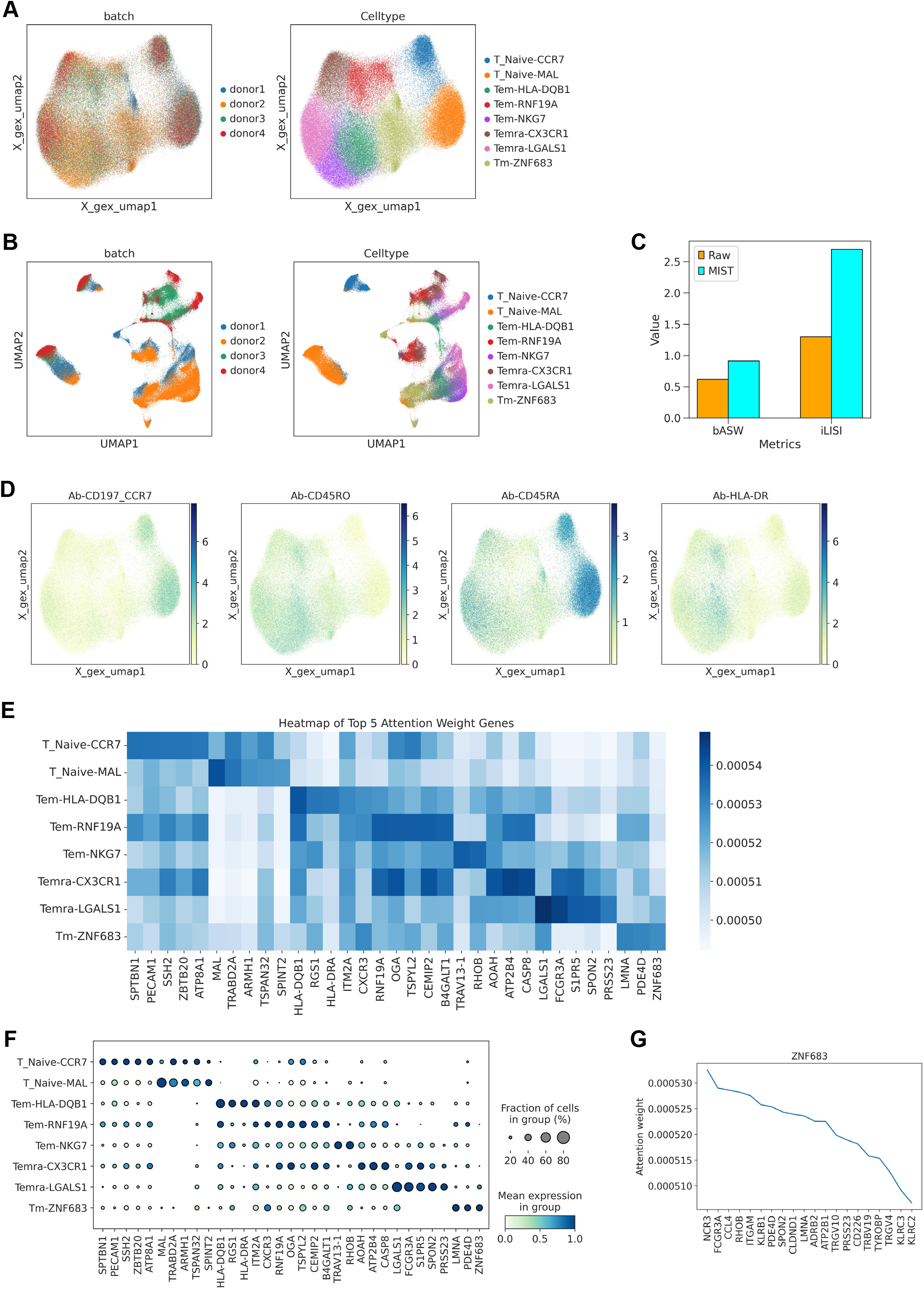
Usage of the GEX latent space for scRNA-seq data analysis. A. UMAP (Uniform Manifold Approximation and Projection) plots of T cells in the 10x_200k dataset by the GEX representation of latent space from MIST, color-coded by donor origin (left) and cell type (right), respectively. Cell types are designated as canonical T cell classification followed by a secondary cell cluster specific gene. T_Naive: native T cell; Tem: effector memory T cell; Temra: terminal effector memory T cell; Tm: memory T cell. B. UMAP plots of T cells in the 10x_200k dataset by raw expression, color-coded by donor origin (left) and cell type (right), respectively. In the left UMAP plot, cells from the same donor are concentrated; in the right UMAP plot, considerable number of cells annotated as distinct types of T cells are mixed together. These observations indicate possible batch effect underlying the dataset. C. Statistical comparison of cell cluster robustness for the 10x_200k dataset when raw expression or GEX latent space from MIST were used for clustering. bASW: batch Average Silhouette Width; iLISI: integration Local Inverse Simpson’s Index. D. Same as A, but with T cell surface protein (CCR7, CD45RO, CD45RA and HLA-DR) expression level mapping. E. Heatmap showing gene-to-cell type attention weight for the top 5 genes per cell type. F. Dot plot for the top 5 genes by attention weight for each cell type. Color scale indicates the mean of normalized expression of genes in each cell type, and dot size is proportional to the percentage of cells within each cell cluster expressing the genes. The same applies to all dot plots in this paper. G. Exemplary gene-wise attention weight. The blue line shows the attention weight of the top 20 genes with highest attention for ZNF683, ranked by attention weight.

We next sought to understand whether the self-attention layer of GEX encoder can provide biologically meaningful insight for the analyzed single-cell transcriptome data. We found that the genes with high attention weights for a cell cluster had highly specific expression among cells in that cluster (Figure 2E-F; Figure S2B). Thus, instead of identifying cell cluster-specifically expressed genes as markers, the gene-to-cell attention weights could be used as the basis for annotating cell clusters. In addition, the genes with high gene-to-gene attention weights were of significantly correlated expression (*9, 22*) (Figure 2G), suggesting the self-attention layer embedded co-expression relationship between genes. Overall, these results derived from the GEX latent space indicate our MIST framework is capable of transforming single-cell transcriptome profiles as composite features with biologically meaningful interpretability, for relevant downstream analyses.

### Interpretation of TCR by MIST

TCR sequence could directly influence T cell function, however, it is unclear to what extend T cell function is dictated by its TCR sequence. We sought to evaluate whether MIST can comprehend the semantics behind TCR sequences and how such semantics translate into functional impact. The TCR sequences in the 10x_200k dataset were in the form of paired TCRα-TCRβ chains, with TCRα containing TRAV-CDR3α- TRAJ, and TCRβ encompassing TRBV-CDR3β-TRBJ. The TRAV, TRAJ, TRBV and TRBJ genes were recorded with V/J gene symbols, while the CDR3 sequences were in the form of amino acid sequences (referred as CDR3αaa and CDR3βaa) (*23*). The TCR encoder of our MIST framework took in the TCR sequences in the 10x_200k dataset as input, transformed V/J genes as separate gene embeddings, and encoded CDR3aa sequences into an amino acid embedding together with a position embedding (Figure S1A). Each CDR3aa sequence was treated as a ‘sentence’ with each amino acid considered as a separate ‘word’. The encoded amino acid embeddings trained using the 10x_200k dataset resulted amino acids with similar biochemical properties being placed relatively close to each other in the PCA plot (*24*) (Figure 3A). This indicated that the trained amino acid embeddings could legitimately reflect the properties of amino acids within CDR3 sequences, making such abstract embeddings biologically interpretable.

**Figure 3:**
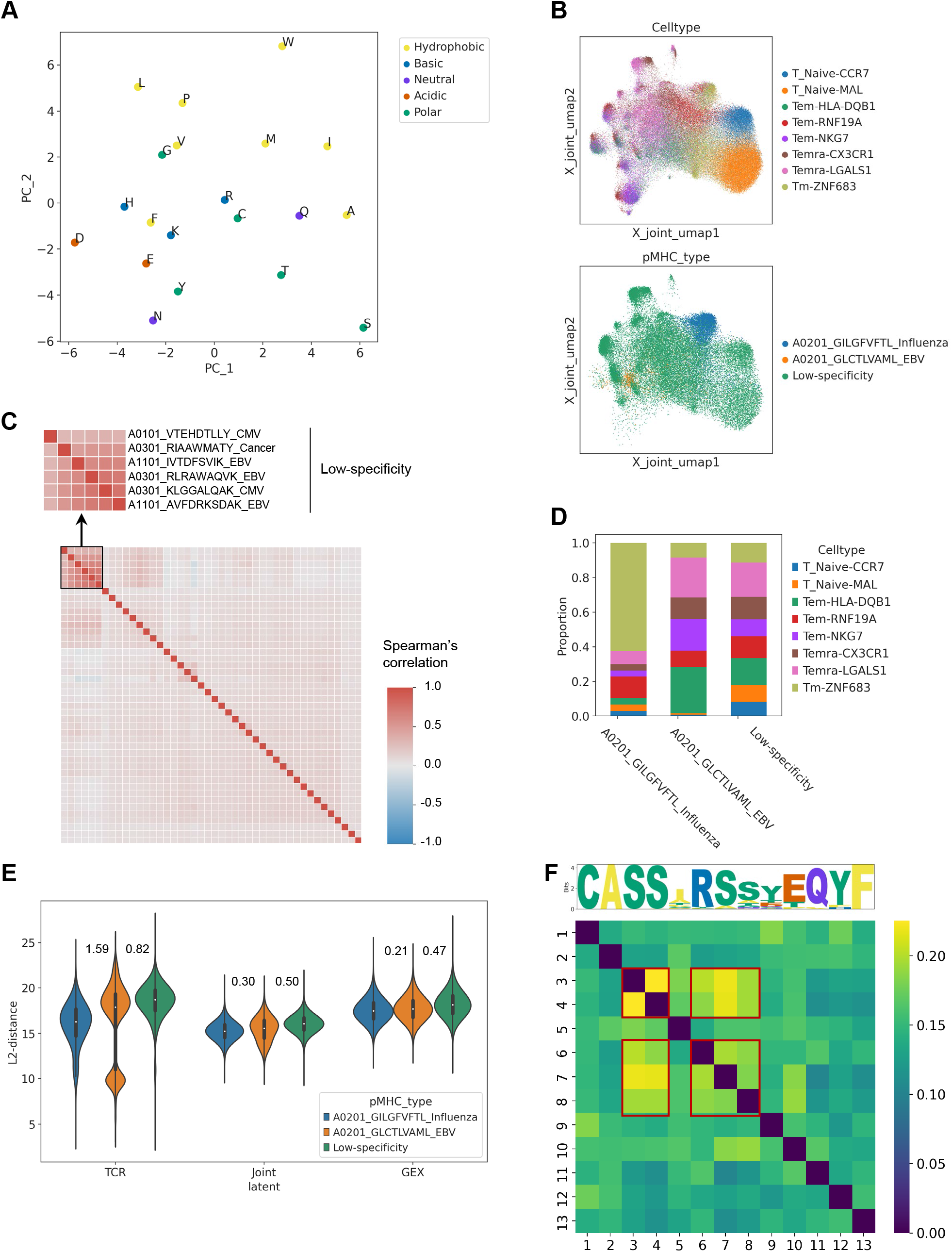
Usage of the joint latent space for combinatorial scRNA-seq and scTCR-seq data analysis for antigen-specific T cells. A. PCA (Principal Component Analysis) plot of 20 amino acids, color-coded by chemical properties. PCAs for the amino acids are derived from the 64-dimensional embedding of each amino acid in TCR CDR3 sequences of the 10x_200k dataset by the TCR encoder in MIST. Refer to Figure S1D and Methods for details. Amino acids with the same chemical property are positioned close to each rather in the PCA plot, indicating similar embedding. B. UMAP plots of T cells in the 10x_200k dataset by the representation of joint latent space from MIST, color-coded by cell type (top) and antigen specificity (bottom). Only T cells with TCR determined as specific to the listed pMHCs in the color-code are shown in the bottom UMAP plot; and the low-specificity pMHCs are listed in Figure 3C. C. Heatmap showing the pairwise Spearman’s correlation between binding affinity to different TCR clonotypes for 44 distinct antigens (pMHCs) in the 10x_200k dataset. The zoom-in region shows 6 antigens with strongly correlated binding profiles, suggesting non-specific or low specific binding to all TCRs by these 6 antigens. Antigens are designated as HLA subtype_epitope peptide_antigen type. D. Proportion of distinct types of T cell with TCR specific to different antigens. E. Violin plots of cell-cell distance (Euclidean) for T cells with distinct types of antigen specificity. Distances are derived from the TCR, GEX and joint latent space representation by MIST, respectively. White circle indicates data median; thick vertical line covers 25^th^ to 75^th^ percentile. The same applies to all violin plots in this paper. The numbers between the adjacent violins represent the absolute value of the difference between medians of the two groups. F. Heatmap showing the pairwise attention weight between amino acids within the CDR3β sequences of TCR specific to influenza. Peptide motif of the CDR3β sequences is shown on top. Only CDR3β sequences with 13 amino acids are included in the analysis. Red boxes highlight conserved amino acids with relatively high pairwise attention.

Similar to the design philosophy of the GEX VAE, the TCR VAE in MIST attempts to maximumly reconstruct the input TCR sequences, and in the process a latent 128D TCR representation is created for each TCR sequence. The principal components derived from the TCR latent space can similarly be used for dimensional reduction and visualization (Figure S2C). Interestingly, the UMAP plot of the TCR latent representations for the 10x_200k dataset grouped T cells into sparse clusters (Figure S2C left), with several clusters corresponding to the same functional annotation by transcriptome profile (Figure S2C right). This is potentially linked to the fact that TCR sequences are abstract and more variable in comparison to the transcriptome data, and the T cells with the same annotation by transcriptome could be of distinct antigen specificity dictated by TCR (*25, 26*).

### MIST incorporates multi-dimensional T cell features

In addition to the GEX and TCR representations, a joint latent space (128D) co-considering both of these representations were computed for each T cell in the 10x_200k dataset (Figure S1A; Methods). UMAP plot derived from this joint latent space surprisingly showed that the naive T cell clusters were still intact as was in the UMAP from GEX representations, however, the memory T cell clusters seemed reformed into new clustering structure (Figure 3B top; Figure S2D). To delve into this, we examined the antigen specificity data of these T cells. Besides the scRNA-seq and scTCR-seq data, the 10x_200k dataset also included quantitative binding affinity information to 44 distinct pMHCs (peptide major histocompatibility complexes) for each assayed T cell (Figure 3C). Projection of these antigen binding affinity data onto the UMAP from the joint latent space showed that T cells clustered closely were of binding specificity to the same pMHC (e.g., A0201_GILGFVFTL_Influenza and A0201_GLCTLVAML_EBV in Figure S3, S4). Further, T cells specific to A0201_GILGFVFTL_Influenza and A0201_GLCTLVAML_EBV were of enrichment for distinct types of memory T cells (*9*) (Tm-ZNF683 and Tem-HLA-DQB1, respectively), comparing to the low-specificity background (Figure 3D). Moreover, we calculated the distances between the T cells with different antigen specificity using the GEX, TCR and joint representations, respectively, and the results showed that 1) T cells with strong antigen specificity were of smaller cell-to-cell distance comparing to the low-specificity T cells, regardless of which representations were used for distance calculation, especially for T cells specific to A0201_GILGFVFTL_Influenza, which is a strong antigen (Figure S5A-B); and 2) the cell-to-cell distances of TCRs with different specificities differ significantly, being most pronounced in the TCR representations, followed by the joint representations, and least significant in the GEX representations, while the joint representations are more stable, and the latent space of TCR are more dispersed (Figure 3B bottom, 3E). These results potentially imply that the joint latent space in MIST could more effectively model cell-to-cell similarity between T cells by combining transcriptome and TCR antigen specificity features.

When we mapped the sample (donor) origin information onto the UMAP by the joint latent space, we found that, once again, the naive T cells from different donors were clustered together without any batch effect, but the memory T cells showed a donor-specific clustering structure (Figure S2D; Figure S5A). For example, clusters of T cells specific to A0201_GILGFVFTL_Influenza pMHC were primarily identified in donors 1 and 2, but not in donor 3 and 4. (Figure S5A). This can be explained by the fact that only donors 1 and 2 were of MHC I subtype of HLA-A0201, thus explicitly having memory T cells recognizing A0201_GILGFVFTL_Influenza. These results reassured that the joint representation from MIST encode composite transcriptome and antigen specificity features for convenient T cell characterization, in both the functional and antigen specificity aspect.

Lastly, the conserved positions in the CDR3 sequence motif of TCRs with the same antigen specificity were of higher inter-position attention weights (*27*) (Figure 3F; Figure S5C-D), and such observation provided biologically relevant interpretation for the attention weights and TCR latent representations.

### Latent representation of T cells in NSCLC using MIST

To further evaluate the capability of MIST, we used it for analyzing the T cells in NSCLC (non-small-cell lung cancer) tumors from patients receiving anti-PD-1 therapy (*28*). The analyzed CD8^+^ T cells (31,066 single cells) were extracted from the scRNA-seq and scTCR-seq data of pre- and post-treatment tumor biopsies (n=46) from 36 NSCLC patients, spanning 32 treatment-naive (TN), 9 site-matched post-treatment (post-rx) responsive (from 8 tumors), and 5 post-treatment non-responsive tumor specimens following combination therapies of PD-1 blockade with chemotherapy. The TN samples included 8 pre-treatment (pre-rx) biopsies from responsive tumors. Once again, GEX encoder of MIST effectively removed batch effect in scRNA-seq data (Figure S6A-C).

We used the joint latent space for the T cells in the NSCLC anti-PD-1 dataset for downstream dimensional reduction and clustering, and the T cells were grouped into 12 clusters, with most of these clusters containing majority of the cells placed close to each other, but cluster 5 to 9 spun out from the majority (Figure 4A left; Figure S6D). When only GEX or TCR latent space were used, very different dimensional reduction patterns were produced attributed to distinct T cell phenotypes and TCR clonotypes (Figure 4A right, 4B; Figure S6E). Albeit the dimensional reduction patterns from the joint and GEX representations were alike for majority proportion of the analyzed T cells, the joint representations were more similar to the TCR representations (Figure 4C), implying TCR clonotypes were of stronger contribution for the joint latent space. Further exploration showed that the GEX and TCR representations for most of the analyzed T cells were independent, except for cells in cluster 5, 7, 8 and 9 (Figure 4D-E; Figure S6F-H), potentially implying T cell function is strongly dictated by TCR clonotype in specific biological environment (*25, 26*).

**Figure 4:**
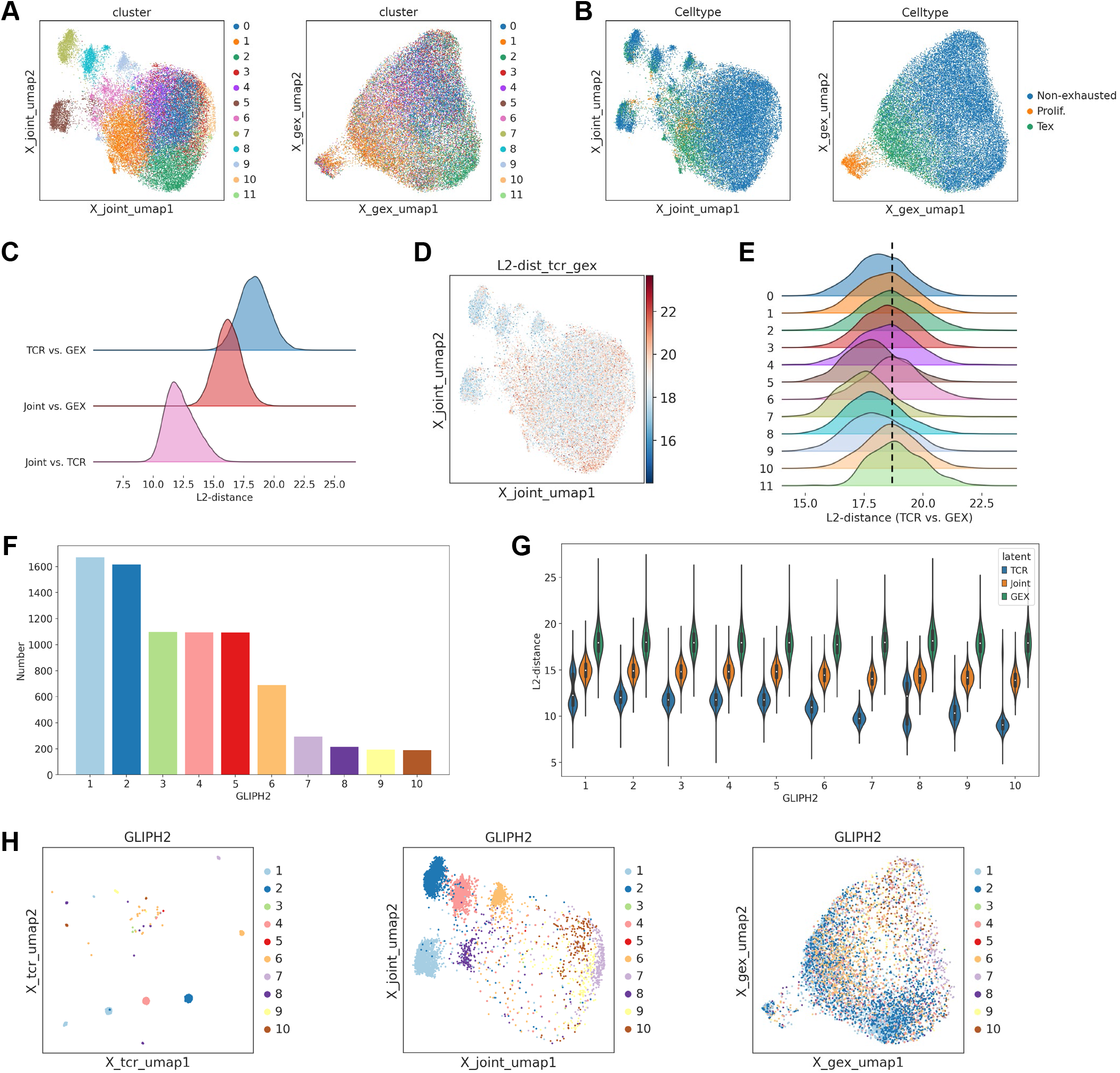
Usage of the latent space for characterization of T cell from NSCLC patients with anti- PD-1 therapy. A. UMAP plots of T cells in the NSCLC anti-PD-1 therapy dataset by the joint (left) and GEX (right) latent space representations from MIST, respectively. Cells are color-coded by cluster ID when clustered using the joint latent space representation. B. Same as A, but cells are now color-coded by T cell phenotype. Tex: exhausted T cell. C. Density plots showing the distribution of Euclidean distances between the latent representations of the GEX and TCR (top), joint and GEX (mid), and joint and TCR (bottom) for each T cells in the NSCLC anti-PD-1 therapy dataset. D. UMAP plot of T cells in the NSCLC anti-PD-1 therapy dataset by the joint representation from MIST, with color reflecting the distance between the GEX and TCR latent representation of each cell. E. Density plots showing the distribution of Euclidean distances between the latent representation of GEX and TCR for T cells in distinct clusters defined in A. F. Bar plot showing the number of T cells (TCRs) in the top 10 clusters identified by GLIPH2 algorithm. GLIPH2 cluster TCRs that are predicted to bind the same antigen. Clusters are ranked according to the number of T cells. G. Violin plots of cell-cell distance (Euclidean) among T cells within each GLIPH2 defined top 10 clusters. Distances are derived from the TCR, GEX and joint latent spaces of MIST, respectively. H. UMAP plot of T cells in the top 10 clusters from GLIPH2 by the TCR (left), joint (mid) and GEX (right) latent representations from MIST, respectively. Cells are color-coded by cluster ID from GLIPH2.

To independently verify the findings from MIST, we used GLIPH2 (*29*) to identify T cell clusters predicted to have similar antigen specificity within the NSCLC anti-PD-1 dataset. We focused on the top 10 clusters identified by GLIPH2 (Figure 4F). Interestingly, the TCR representation produced the smallest distances between cells within each of these GLIPH2 defined clusters, rather than using the joint nor GEX representations (Figure 4G-H). This is consistent with the fact that GLIPH2 only uses TCR sequences for analysis. Importantly, the GLIPH2 defined clusters had strong agreement with the clustering patterns derived using both the TCR and joint latent space from MIST (Figure 4H left and mid), which once again prove that MIST can effectively resolve and capture antigen specificity in TCR sequences.

### MIST identifies T cells responsive to anti-PD-1

We further explored the biological significance for the T cell clusters derived from the joint latent space. We mapped the sample origin, treatment condition and outcome information onto the joint latent representations derived UMAP plot, and then it was clear that cells in clusters 5-9 were almost exclusively from the tumors responsive to anti-PD-1 therapy (Figure S7A). In contrast, such responsive/non-responsive condition separation is not readily visible in the UMAP derived from the GEX representations (Figure S7B). Then we projected the expression level of selected genes important for T cell function and response to anti-PD-1, onto the joint latent representations derived UMAP (Figure S7C). It was clear that the anti-PD-1 responsive T cells (in clusters 5-9) were of high expression of CXCL13 and PDCD1 (encoding PD-1 receptor) (Figure S7C). CXCL13^+^ T cells were reported to better respond to anti-PD-1 (*28, 30, 31*).

Besides clusters 5-9, T cells in cluster 1 also had high expression of CXCL13 and PDCD1. Thus, we sought to understand the biological reasoning for separating these CXCL13^+^ T cells in cluster 1 from clusters 5-9 by using the joint latent representations. Since cells in clusters 5-9 were found in both the pre- and post-treatment tumors that eventually responded to anti-PD-1, we split these cells according to the treatment condition, and refer them as CXCL13^+^-pre and CXCL13^+^-post T cells, respectively. We first identified the genes with strong gene-to-cell attention weight by MIST for cells in cluster 1, CXCL13^+^-pre and CXCL13^+^-post, and confirmed these genes indeed had high average expression in respective cell cluster (Figure 5A-B). Further, T cell functional characteristics scoring showed that cells in cluster 1 and CXCL13^+^-pre were significantly more exhausted and stressed, and in contrast, CXCL13^+^-post T cells were more effector memory-like (Figure 5C-D). Moreover, T cells in cluster 1 had high expression of TIGIT, FOXP3 and IL2RA (Figure 5E; Figure S7D), implying these cells are potentially of regulatory role (i.e., CD8^+^ Treg-like) (*32, 33*), and were functional different from the CXCL13^+^-pre and CXCL13^+^-post T cells. These findings suggest the CXCL13^+^-post T cells were transited from CXCL13^+^-pre by anti-PD-1, rather than cells in cluster 1, despite also expressing CXCL13 (*34*). Such insights were not reported by the original study (*28*). Therefore, in addition to CXCL13, other markers need to be considered to accurately determine the response of T cells to anti-PD-1 therapy. Lastly, MIST could easily identify T cells potentially responsive to anti-PD-1 in the peripheral blood of NSCLC patient (Figure 5F), suggesting MIST could be used for predicting immunotherapy prognosis based on the TCR latent space (*35*).

**Figure 5:**
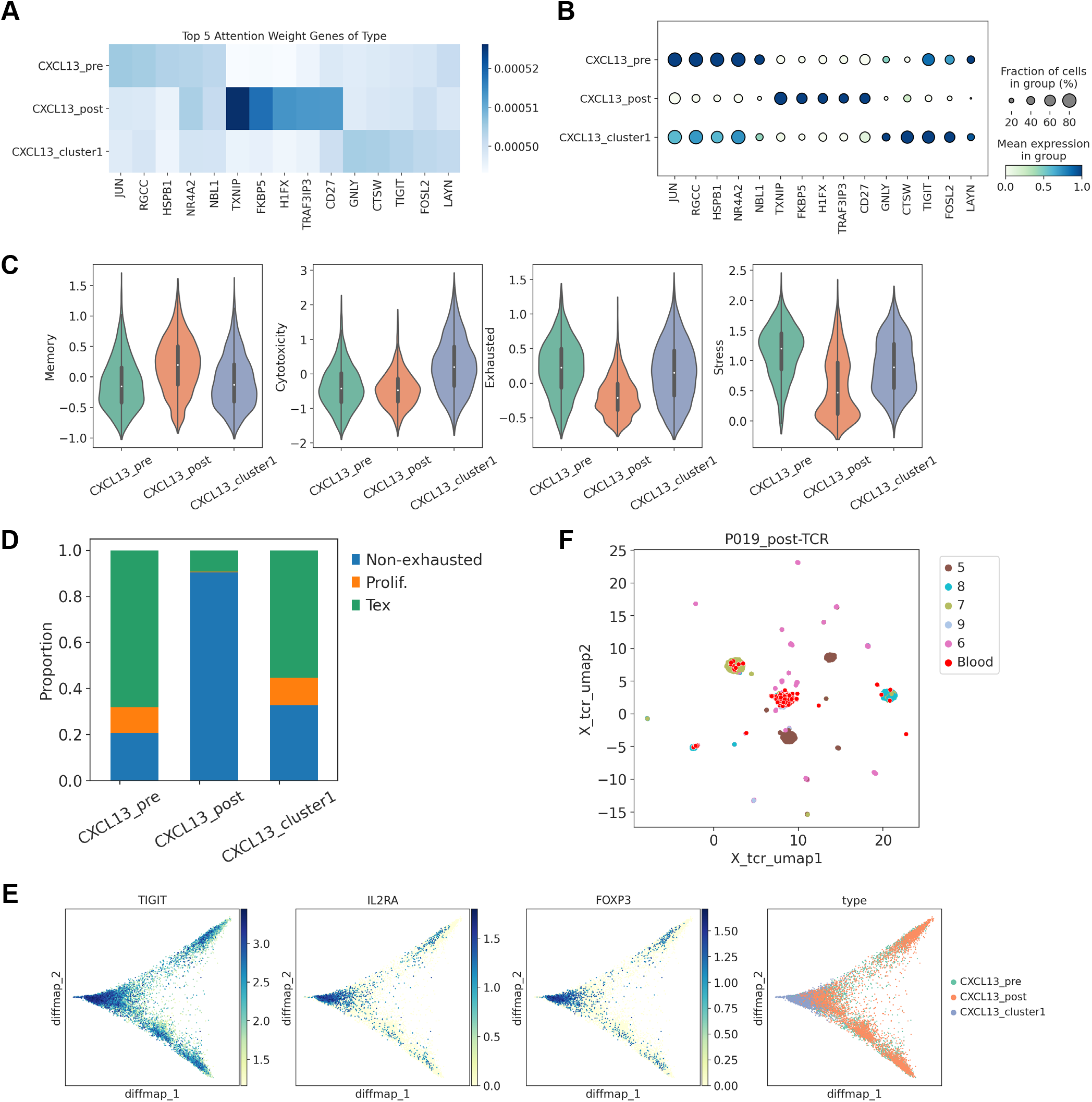
Discovery and prediction of anti-PD-1 therapy responsive T cells by MIST. A. Heatmap showing gene-to-cell cluster attention weight for the top 5 genes per cluster. B. Dot plot for the top 5 genes per cluster by attention weight. C. Violin plots of T cell functional score (memory, cytotoxicity, exhaustion, and stress, respectively) for cells in distinct CXCL13^+^ T cell clusters. D. Proportion of T cells with distinct phenotypes in different CXCL13^+^ T cell clusters. E. From left to right: diffusion map visualization of CXCL13^+^ T cells, color-coded by imputed expression level of TIGIT, IL2RAand FOXP3, and cell type, respectively. F. UMAP plot of T cells from patient P019 after anti-PD-1 therapy by the TCR latent representation from MIST, color-coded by cluster ID as in Figure 4A. T cells drawn as red dot are from peripheral blood sample of P019.

## Discussion

MIST is a deep learning model specifically designed for the integrative analysis of scRNA-seq and scTCR-seq data from T cells. MIST leverages joint representations to learn a comprehensive multimodal model of T cells, effectively linking GEX and TCR features. The joint latent space from MIST is biologically informative for T cell transcriptomes and antigen specificity. In addition to integrating GEX and TCR features, MIST provides numerous essential functions for single-cell sequencing data analysis. These functions are made possible by MIST’s modular design, which projects GEX and TCR into their respective latent spaces.

Additionally, MIST incorporates word embedding and self-attention mechanisms (*13, 14, 22*), techniques commonly applied in natural language processing (NLP). Those approach enhances MIST’s interpretability and allows for detailed specificity analysis of T cell. For transcriptome, MIST can identify co-expression relationships between genes and highlight cell-specific gene expression patterns, offering an alternative to traditional differential gene expression analysis. For TCR, MIST can identify potential key amino acid motifs related to antigen specificity. These capabilities highlight MIST’s flexibility and broad applicability across various scenarios, including the exploration of antigen-specific T cells and tumor immunotherapy-specific T cells, setting it apart from previous algorithms and tools.

MIST stands out in the landscape of T cell analysis tools by offering a distinctive combination of flexibility, interpretability, and robustness. Unlike CoNGA (*9*) and Tessa (*10*), which assume that all T cells with identical TCR clonotypes exhibit similar transcriptome profiles, MIST leverages a deep learning VAE framework to project T cell multimodal features. This approach makes MIST more versatile across diverse biological contexts. Additionally, MIST adopts a holistic strategy by integrating the features of GEX and TCR into a unified latent space, enabling a comprehensive analysis that considers both transcriptome and antigen specificity. In contrast to scNAT (*11*), which merges scRNA-seq and scTCR-seq data into a single latent representation, MIST’s modular architecture maintains separate latent spaces for GEX and TCR. This design enhances its flexibility for downstream analyses, including clustering, GEX imputation, and TCR similarity calculation. Moreover, MIST offers interpretability for both the GEX and TCR features of T cells—a feature that scNAT lacks. Similar to our model, recently published mvTCR (*36*) also provides multiple latent spaces, but MIST’s embedding of TCR takes V/J genes into account, offers interpretability for GEX, and learns the latent relationships between GEX and TCR. Additionally, MIST eliminates batch effects in scRNA-seq data through the DSBN layer, a consideration that has been largely ignored in most previous models (*10, 11, 36*).

MIST holds promise for expansion beyond its current capabilities. Its architecture is potentially adaptable for the integration of B cell receptor (BCR) data, γδ T cell receptor data, and other omics datasets such as single-cell ATAC-seq (scATAC-seq). For B cells and γδ T cells, their adaptive immune receptors share similar serialized representations with αβ T cells (*3, 4, 37*), making them suitable for MIST’s framework. Additionally, while obtaining scATAC-seq and scTCR-seq data from the same T cell barcode is currently not feasible, the success of similar modular models like MultiVI (*15*) in integrating scATAC-seq and scRNA-seq suggests that MIST could potentially be adapted to integrate scATAC-seq, scRNA-seq, and scTCR-seq in the future. This adaptability could significantly enhance its utility in multi-omic studies, providing a more comprehensive understanding of immune cell dynamics.

Despite its strengths, MIST does have some limitations. One notable drawback is its inability to generate data from one modality based on another, meaning it cannot generate TCR from GEX or vice versa. This limitation arises from our approach to modeling the relationship between GEX and TCR, which avoids overfitting the two modalities to maintain their distinct characteristics and representations. To better utilize single-modality data, we employed a batch-independent encoder trick (*18*). Additionally, the training time for MIST is relatively longer compared to training models on individual scRNA-seq or scTCR-seq data alone, due to the complexity of integrating multimodal data.

In sum, MIST provides a flexible, interpretable, and robust framework for analyzing T cells at single-cell resolution. By offering detailed insights into GEX and TCR, MIST opens new avenues for understanding T cell biology and its role in health and disease. This comprehensive approach ensures that MIST is a powerful tool for researchers aiming to explore the complexities of the immune system at single-cell resolution.

## Methods

### MIST architecture

MIST utilizes a modular VAE to project GEX and TCR data into latent space. As a result, GEX and TCR data assign their own latent representations, modeled as isotropic multivariate normal distributions. To unify these representations, the latent representations are averaged 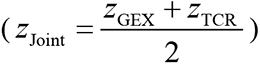 as MultiVI (*15*) (Figure S1A). MIST also incorporates a weighted scheme for GEX and TCR representations, utilizing learnable weights. This joint representation is then used to decode all model parameters.

The GEX encoder is a three-layer neural network (self-attention layer – fully connected layer – batch normalization (BN) – ReLU – fully connected layer) that calculates the mean *μ_GEX_* and variance *σ* ^2^*_GEX_* of the 128-dimensional latent space, utilizing reparameterization to obtain the latent representation *z_GEX_* . To manage the computational complexity of long sequences (2000 gene inputs), linearly scalable attention (Performer) (*14*) is employed to process the GEX data.

The attention matrix (Performer) is described as follows:

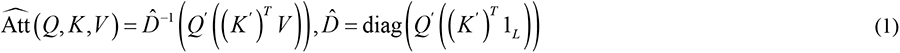

whlie *Q*^′^ = ∅(*Q*), *K* ^′^ = ∅(*K*), and ∅(*x*) is defined as:

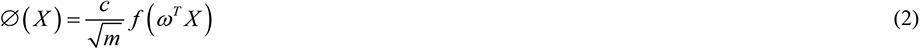

where *c* is a positive constant, *ω* is a random feature matrix, and *m* is the dimesionality of the matrix.

The GEX decoder is a two-layer neural network (average pooling layer – fully connected layer – DSBN – Sigmoid) for reconstructing GEX data, with an average pooling layer designed to prevent overfitting. The DSBN layer utilizes different BN layers for different data batches, enabling the model to transform domain-specific data into domain-invariant representations, effectively removing batch effects (*17, 18*). By placing DSBN in the decoding layer, the encoder becomes batch-independent, allowing it to project new data without requiring retraining.

The reconstruction loss function for the GEX is the binary cross-entropy between the input data and the reconstructed data:

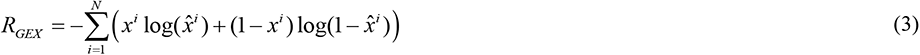

The TCR encoder first embeds the V/J genes and CDR3aa sequences into learnable vectors, with embedding dimensions of 48 for genes and 64 for each amino acid. The features of CDR3aa sequences is then concated with the gene features after passing through a Transformer-like self-attention module and a three-layer convolutional neural network (with 16, 32, and 64 output channels, respectively) (*13, 38*).

Finally, the mean *μ_TCR_* and variance *σ*^2^*_TCR_* of the 128-dimensional GEX latent space are obtained through two fully connected layers, and the latent representation *z_TCR_* is derived through reparameterization.

Since the CDR3aa sequences is short, with most sequences being within 30 amino acids, regular dot-product attention is used in the self-attention module.

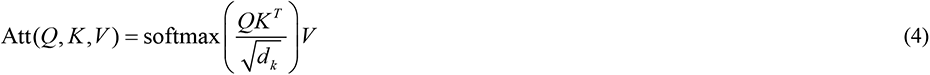

The TCR decoder reconstructs the CDR3aa sequences and the V/J gene separately. For the CDR3aa sequences, it passes through a fully connected layer and two layers of de-convolutional neural networks (with 32 and 64 output channels, respectively), and finally through another fully connected layer. The gene reconstruction primarily involves two fully connected layers. All final fully connected layers in the TCR decoder share weights with their corresponding embedding layers. The reconstruction loss of the TCR is calculated using cross entropy. Based on prior biological knowledge, the J gene segment of TCR has a low impact on the specific antigen recognition function of TCR (*39, 40*), so its loss is weighted by 0.1.

Reconstruction loss function of TCR single chain is defined as:

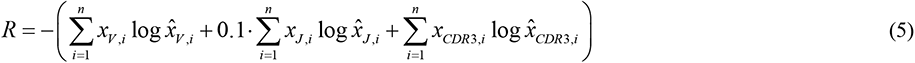

Reconstruction loss function of TCRα and TCRβ chain is thus defined as:

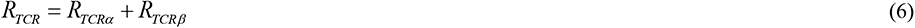

In order to learn the potential connection between GEX and TCR data, we added a penalty term to minimize the distance between their representations. Unlike chromatin accessibility data, from which GEX can be more accurately inferred (*41*), it is almost impossible to accurately infer GEX from TCR sequences. Overfitting the representations of GEX and TCR may result in the loss of critical features in the GEX data. Thus, instead of using the symmetric Jeffrey’s divergence in MultiVI (*15, 42*), we chose maximum mean discrepancy (MMD) (*16*). MMD offers greater flexibility compared to symmetric Jeffrey’s divergence as it does not require the supports of the distributions to overlap and can be adapted through the choice of kernel functions (*43*).

The Gaussian Kernel is defined as:

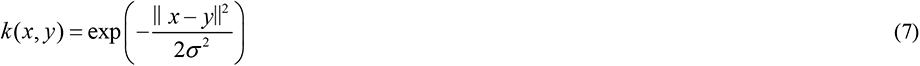

For the sample sets {*z_GEX_* _,*i*_ } and {*z_GEX_* _, *j*_ }, kernel function matrix is defined as:

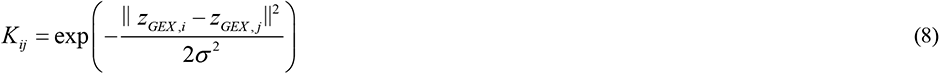

MMD is defined as:

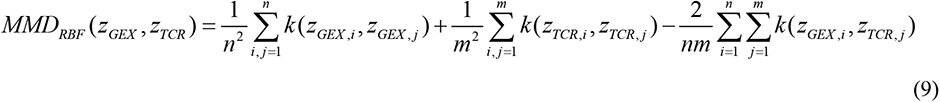

The variational loss of MIST is the KL divergence between the joint representation and the standard Gaussian distribution. In addition, the variational loss weight is set to 0.5 for the more flexible alignment under latent space:

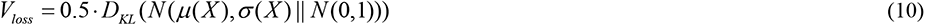

In conclusion, the total loss includes four parts: the GEX reconstruction term, the TCR reconstruction term, the penalty term, and the variational loss, formulated as:

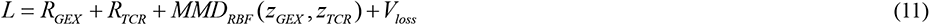

### Training process

MIST is optimized using AdamW (*44*) with a learning rate of 0.0001 and weight decay of 0.001 by default. We split the input data into training and validation sets at a ratio of 0.9 to 0.1. The maximum number of model training epoch is set to 300 by default. If the total loss on the validation dataset does not improve within 30 consecutive epochs (1/10 of the maximum number of epochs), early stopping is applied (*45*). Additionally, we down-weighted the variational loss during the first 30 epochs (1/10 of the maximum number of epochs) to allow the model to fully learn the reconstruction and potential connections of multimodal data 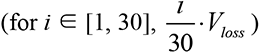 (*15*). The code and training procedures for MIST are based on the PyTorch framework (*46*). Anndata object is computed after training for further analysis and visualization with scanpy (*47*).

### Attention weights for model interpretability and redundant gene removal

In NLP, the attention weights in an attention matrix reflect the contribution and interaction of each token pair (*13*). By averaging the attention matrices across all heads and layers, we can integrate all the attention matrices into a single matrix. In this averaged attention matrix, each value represents the degree of attention that token *i* pays to token *j*. For MIST’s attention matrix, this corresponds to the attention that gene *i* pays to gene *j*, or that amino-acid *i* pays to amino-acid *j*. To focus on the importance of genes to T cells or amino acids to TCR, we average the columns of the attention matrix to form an attention vector, with its length equal to that of the sequence. This approach allows us to identify top attention genes for specific cell types. Additionally, since GEX may contain redundant genes that have high counts across most cells, leading to high attention weights, we propose a redundancy removal function. This function includes two parameters: ‘min_occurrence’ and ‘top’. If a gene appears in the ‘top’ attention weight genes by a given cell proportion (‘min_occurrence’), it is considered redundant and is removed.

### Modeling differences between MIST and scNAT

While conceptually similar, MIST and scNAT (*11*) have several key differences in their design and implementation choices. We detail several distinctions between MIST and scNAT: (1) MIST employs a distributional averaging and penalization approach to merge latent representations, whereas scNAT directly concatenates them. This allows MIST to include multiple latent representations. (2) MIST incorporates a self-attention mechanism to enhance interpretability, whereas scNAT lacks interpretability. (3) MIST utilizes pooling, an early stopping strategy, and dropout techniques to prevent model overfitting and enhance its generalizability. (4) scNAT requires additional model processing for GEX input, whereas MIST only requires log normalized expression counts as input.

### scRNA-seq and scTCR-seq data preprocessing

The scRNA-seq data were imported and processed using scanpy (*47*). Cells with fewer than 600 detected genes were excluded, and only genes expressed in more than 3 cells were retained. The GEX were log-normalized with a size factor of 10,000, while the membrane protein expression values were centralized using log-ratio normalization. Subsequently, 2000 top variable genes specific to the batch were selected. Values of each gene were normalized to a range of 0 to 1 using the MaxAbsScaler function from the scikit-learn package (*48*).

The scTCR-seq data were imported and quality-controlled using the scirpy (*49*). Only T cells with complete α and β chains were retained, and TCR with CDR3aa sequences longer than 30 amino acids were excluded. Dictionaries were created for amino acids and each of the V/J genes, converting the amino acids and genes into vectors. To ensure consistent model input, CDR3aa sequences were zero-padded to a length of 30 as typically done in NLP. Finally, the processed scRNA-seq GEX matrix and scTCR-seq data were used as input for MIST.

### Dimensionality reduction and clustering

Using PCA from the scikit-learn package (*48*), the 128-dimensional latent representations (GEX, TCR and joint, respectively) were reduced to 15 dimensions. Subsequently, scanpy’s ‘pp.neighbors’ function was used to construct the graph, followed by clustering with the ‘tl.leiden’ function and non-linear dimensionality reduction with the ‘tl.umap’ function. For the top variable gene matrix of normalized expression counts, PCA was directly applied for dimensionality reduction, serving as a batch effect control.

### Batch correction metrics

#### Batch average silhouette width (bASW)

The silhouette score measures the relationship between the within-cluster distance of a cell and the between-cluster distance of that cell to the closest cluster (*19*). The batch ASW is computed based on the absolute silhouette of batch labels for each cell *i* of cell type *j*, then averaged across *M* cell types (*50*).

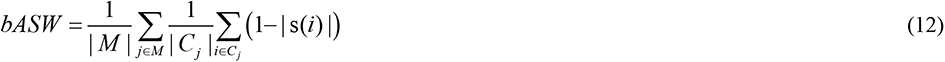

Batch ASW values range from 0 to 1, with 0 indicating strongly separated batches and 1 indicating ideal mixing of batch.

#### Integration local inverse Simpsons Index (iLISI)

We calculated iLISI values using the ‘compute_lisi’ function in the lisi R package (*20*). UMAP embeddings and batch labels were used as input in calculation.

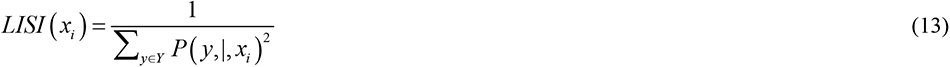

where *x_i_* ∈{*x*_1_, *x*_2_, …, *x_N_*} is the *i*-th cell’s UMAP embeddings in the dataset of size *N*, and *Y* is the set of batch label values with respect to the type of LISI we are computing. The probability *P* ( *y*,|, *x_i_*) refers to the ‘relative abundance’ of the covariate y within KNN (k-nearest neighborhood) of *x_i_* . iLISI values are greater than 0, with larger values indicating more ideal mixing of batches.

### L2-distance calculation

The distance between latent representations after PCA dimensionality reduction is calculated using Euclidean distance (L2-distance), which can be categorized into within-cluster and between-cluster distances. The within-cluster L2-distance indicates the dispersion and compactness of each cluster; the more dispersed the cluster, the larger the L2-distance, and vice versa. The between-cluster L2-distance reflects the similarity and proximity between clusters; the more similar the clusters, the smaller the L2-distance, and vice versa.

### GLIPH2 analysis

To identify TCR specificity groups for the NSCLC anti-PD-1 therapy dataset (NCBI GEO accession: GSE179994) (*28*), GLIPH2 analysis was carried out as described previously. GLIPH2 clusters TCRs based on 1) global similarity, calculated by CDR3 sequences differing by up to one amino acid, and 2) local similarity, calculated by shared enriched CDR3 motifs (*29*). The filtering criteria for GLIPH2 clusters were set to groups with significant final score (*P* < 0.00001) and significant V-gene bias (*P* < 0.005) (*11*).

### GEX imputation, functional score and diffusion map

In MIST, GEX imputation bridges the MAGIC algorithm (*51*), where the GEX latent representation is PCA- dimensionally reduced to compute the Markov chain. Additionally, we used the ’tl.score_genes’ function in scanpy to calculate functional gene set scores for each single cell. The memory T cell gene set includes IL7R, GZMK, CD44, CD27, CD28, and CXCR3 (*52, 53*); the cytotoxicity gene set includes GNLY, GZMB, PRF1, NKG7, and IFNG (*54, 55*); the exhaustion gene set comprises HAVCR2, LAG3, TIGIT, CTLA4, PDCD1, CD160, and TOX (*55–57*); and the general stress gene set includes BAG3, CALU, DNAJB1, DUSP1, EGR1, FOS, FOSB, HIF1A, HSP90AA1, HSP90AB1, HSP90B1, HSPA1A, HSPA1B, HSPA6, HSPB1, HSPH1, IER2, JUN, JUNB, NFKBIA, NFKBIZ, RGS2, SLC2A3, SOCS3, UBC, ZFAND2A, ZFP36, and ZFP36L1 (*54, 58*). Finally, for the diffusion map computation, the data was preprocessed by computing a neighborhood graph using 30 neighbors and the first 15 principal components. The first 3 components of the diffusion map were then computed.

## Data and code availability

All data used in this paper are publicly accessible through the original publications and releases. Data for NSCLC anti-PD-1 therapy were download from NCBI GEO, accession number GSE179994. The 10x_200k dataset is available from the website: https://www.10xgenomics.com/resources/datasets. Python source code and MIST are publicly available on GitHub at https://github.com/aapupu/MIST.

## Competing interests

The authors declare that they have no competing interests.

## Authors’ contributions

WP.L. and O.J.L. conceived and developed the model, wrote the code, performed sequencing data analysis, and wrote the manuscript with input from Y.Q.L..

## Acknowledgements

This study was supported by funding from the Natural Science Foundation of China (82350610280 and 92370107); the Pearl River Talents Scheme of Guangdong Province (2019QN01Y990). O.J.L. gratefully acknowledges the support of K. C. Wong Education Foundation.

---

**Figure S1:**
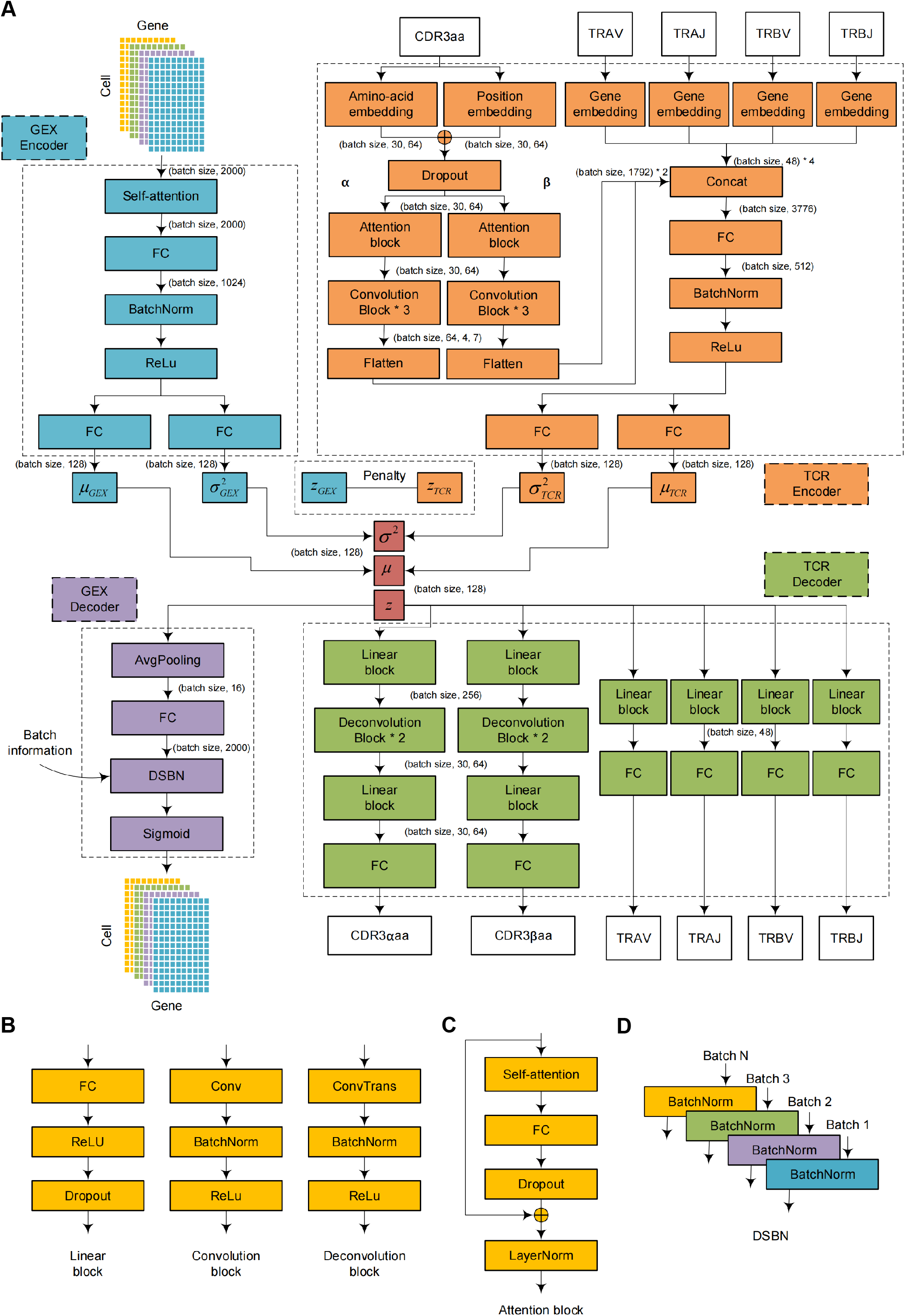
Detailed architecture of the MIST model. A. Diagram showing detailed architecture and workflow of the MIST model. FC: fully connected; BatchNorm: Batch Normalization; DSBN: Domain-Specific Batch Normalization; CDR3aa: TCR CDR3 amino acid sequence; CDR3αaa: CDR3 amino acid sequence of TCRα; CDR3βaa: CDR3 amino acid sequence of TCRβ. Batch size is a user customizable parameter. Numbers in parentheses indicate the dimension of the output for each layer. B. Diagrams showing detailed architecture and workflow of the Linear, Convolution and Deconvolution blocks in the MIST model. Conv: Convolution; ConvTrans: Transposed Convolution. C. Diagram showing detailed architecture and workflow of the Attention block in the MIST model. LayerNorm: Layer Normalization. D. Diagram showing detailed workflow of DSBN.

**Figure S2:**
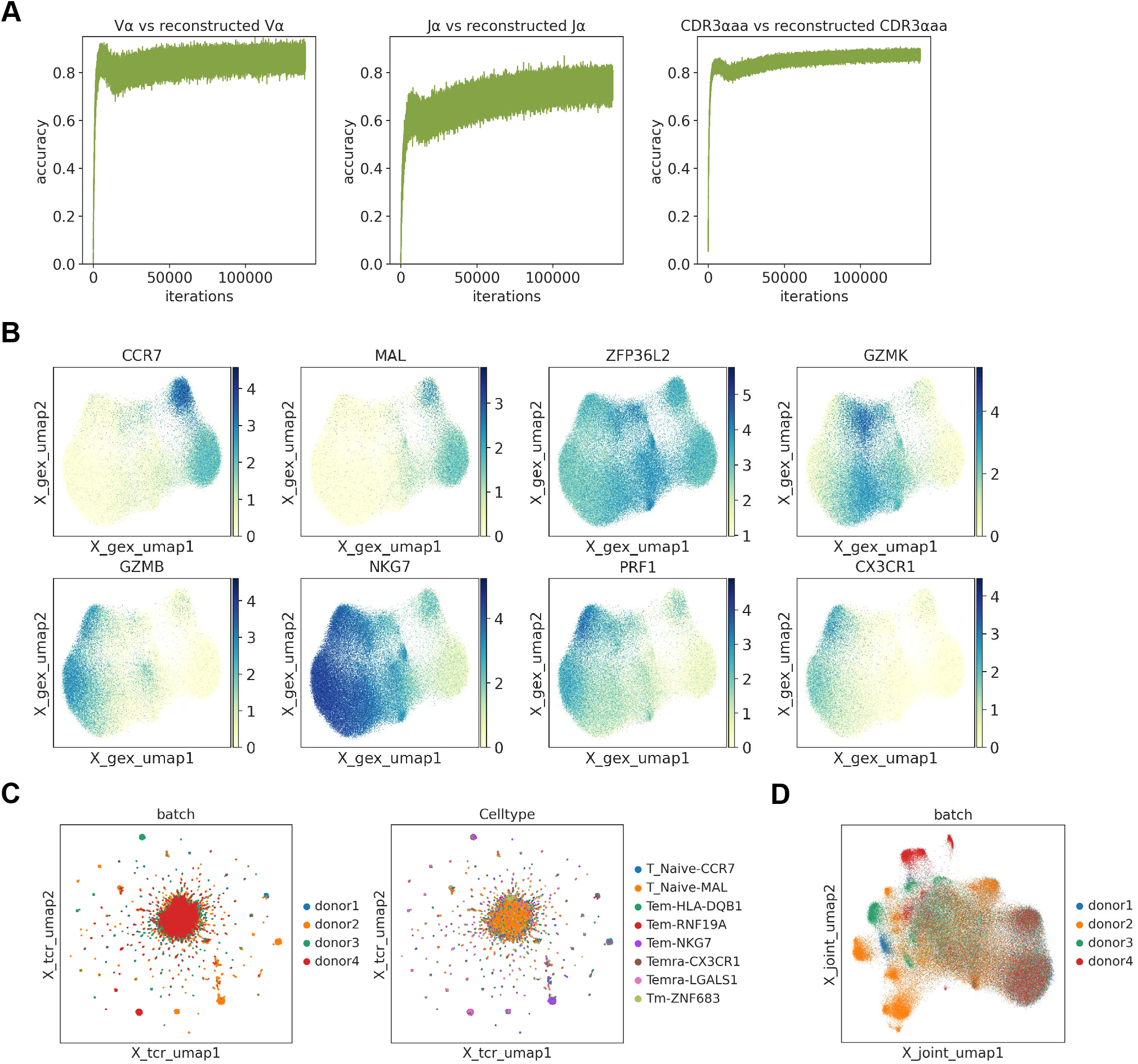
Characterization of T cells in the 10x_200k dataset by the MIST model. A. Line graphs illustrating the increases in accuracy between the input V/J gene (left and mid) and CDR3aa (right) of TCRα and the corresponding reconstructed TCRα segments over training cycles for the 10x_200k dataset. B. UMAP plots of T cells in the 10x_200k dataset by the GEX latent representation from MIST with selected cell type marker gene expression level mapping. C. UMAP plot of T cells in the 10x_200k dataset by the TCR latent representation from MIST, color-coded by donor origin (left) and cell type (right). D. UMAP plot of T cells in the 10x_200k dataset by the joint latent representation from MIST, color-coded by donor origin.

**Figure S3:**
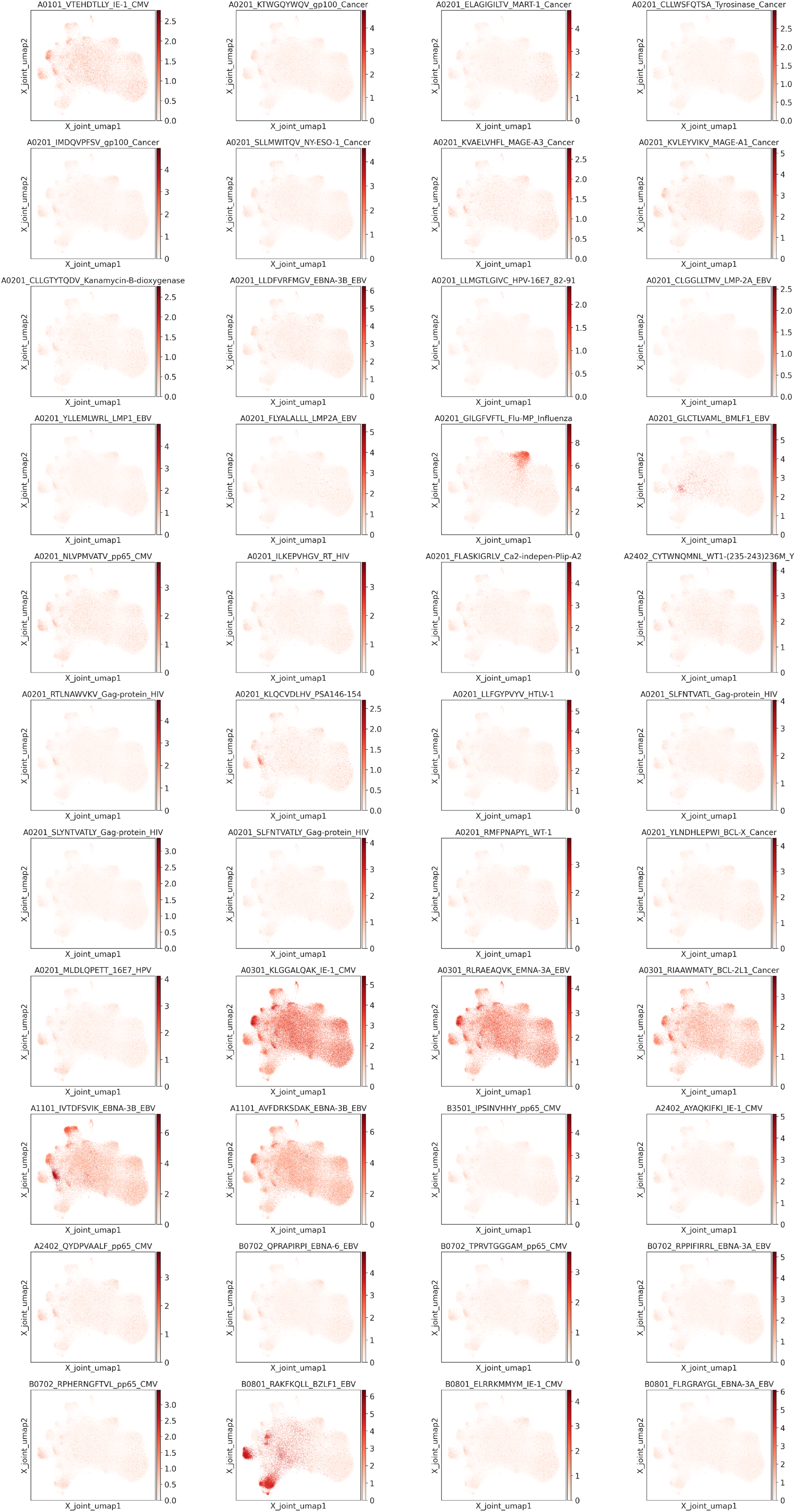
Characterization of antigen binding affinity of T cells in the 10x_200k dataset. UMAP plots of T cells in the 10x_200k dataset by the joint latent representation from MIST, with color reflecting binding affinity to each antigen (pMHC; 44 in total), as shown on top of each plot. The binding affinity is calculated as log(UMI count + 1) per antigen per cell, and UMI count is determined by sequencing of barcoded MHC multimer loaded with antigenic epitope, bound to each T cell. Higher UMI count means higher binding affinity.

**Figure S4:**
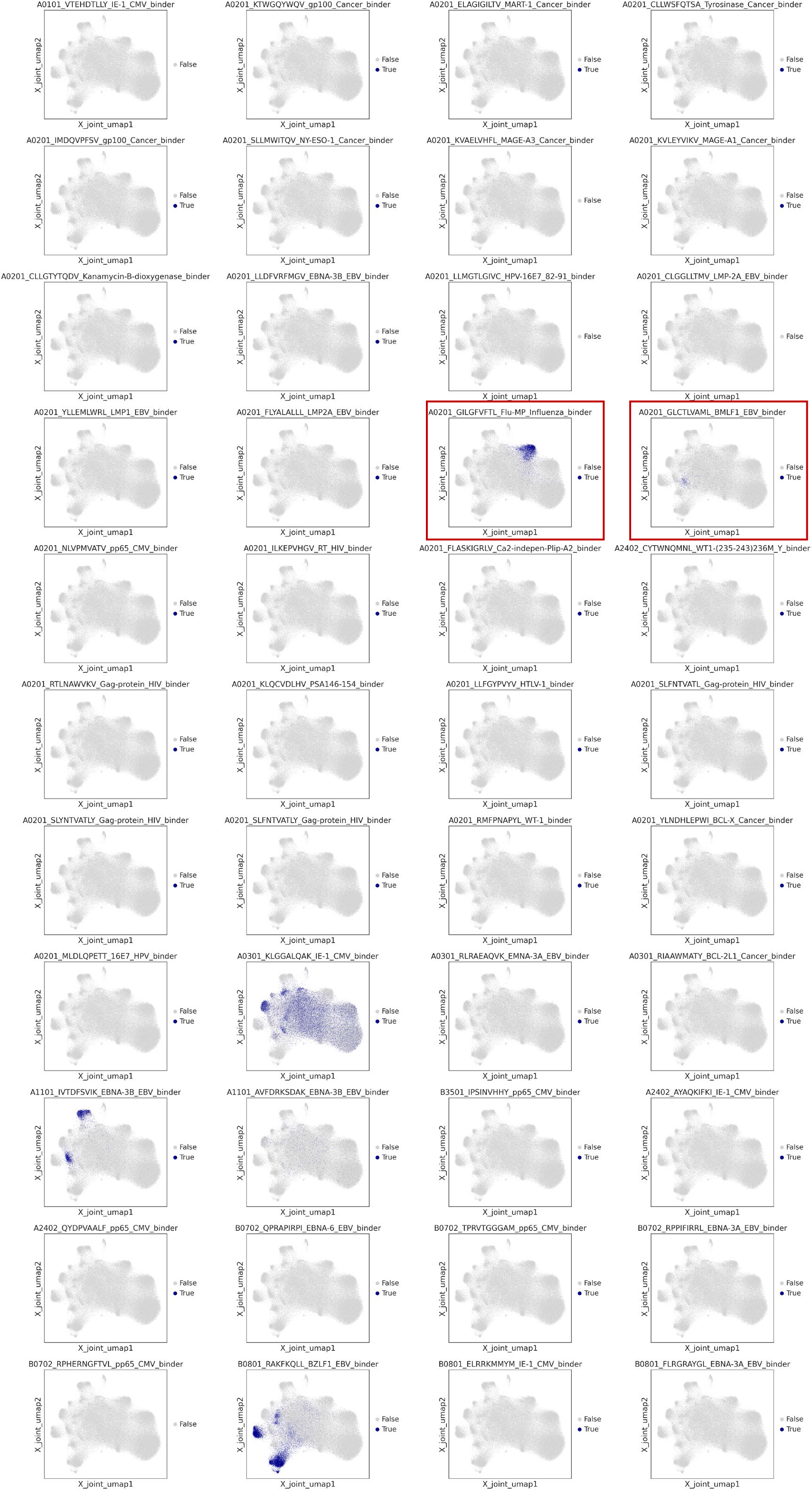
Characterization of antigen specificity of T cells in the 10x_200k dataset. UMAP plots of T cells in the 10x_200k dataset by the joint representation from MIST, with color reflecting antigen (pMHC; 44 in total) specificity status. pMHC is shown on top of each plot. A T cell is determined as specific to an antigen if UMI count greater than 10 and that also greater than five times the highest negative control UMI count for the corresponding pMHC (according to the protocol from 10X Genomics). UMI count is determined by sequencing of barcoded MHC multimer loaded with antigenic epitope, bound to each T cell.

**Figure S5:**
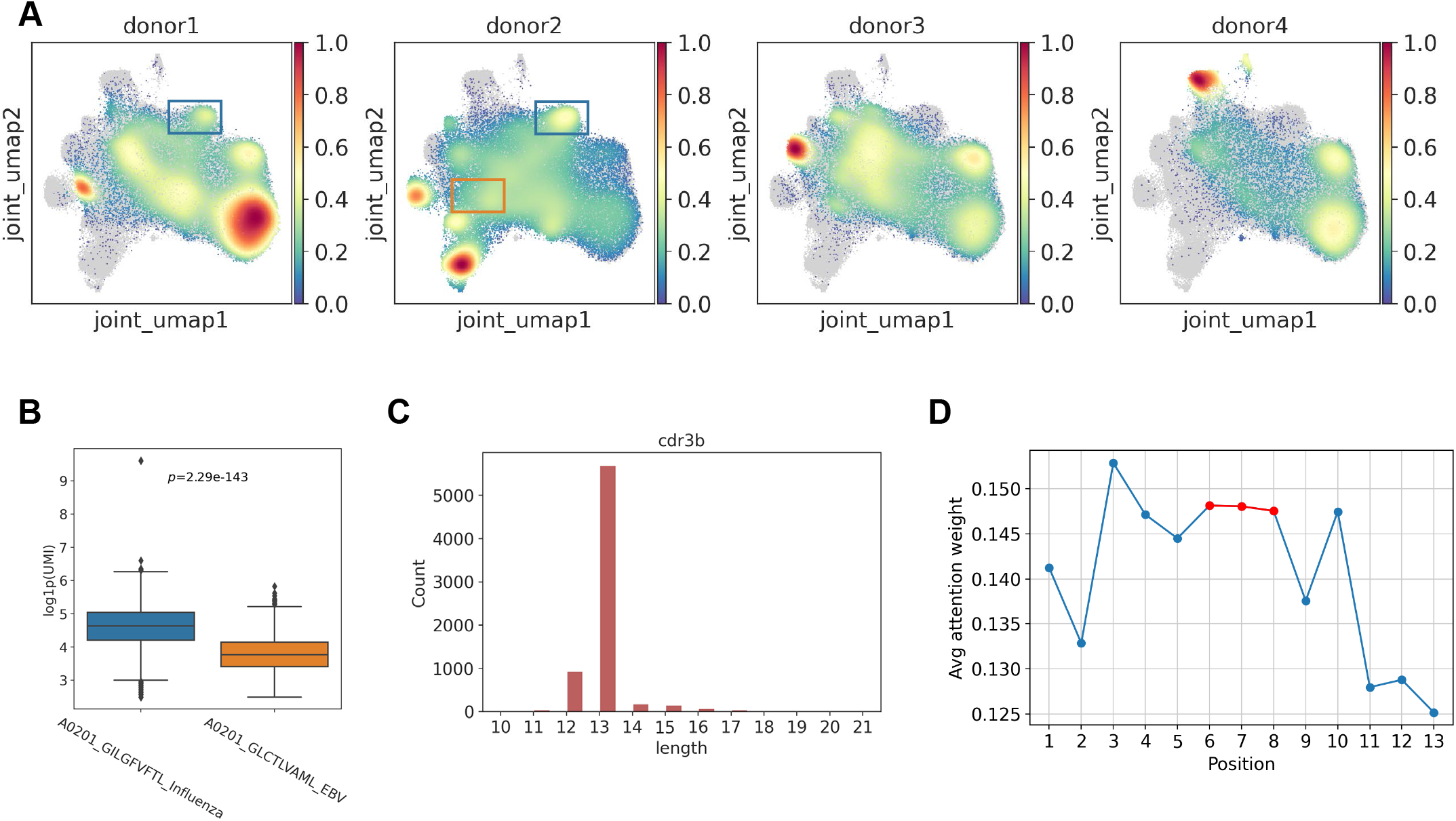
Additional characterization of interpretability of the MIST model. A. UMAP plots of T cells in the 10x_200k dataset by the joint latent space from MIST per donor, with color reflecting density of cells from different donor. The blue and orange boxes highlight T cells with strong binding affinity to A0201_GILGFVFTL_Influenza and A2021_GLCTLVAML_EBV pMHCs, respectively. B. Boxplots of T cell binding affinity to distinct antigens in the 10x_200k dataset. The binding affinity is calculated as log (UMI count + 1) per antigen per cell, and UMI count is determined by sequencing of barcoded MHC multimer loaded with antigenic epitope, bound to each T cell. The outlines of the boxes represent the first and third quartiles, the line inside each box represents the median, and boundaries of the whiskers are found within 1.5 times the interquartile range, with dots representing outliers. *P* value was calculated by Wilcoxon Rank Sum test (two-sided). C. Histogram showing distribution of sequence lengths for influenza specific TCR CDR3β of influenza. D. Line graph showing the average attention weight of each amino acid position within the CDR3β sequences of TCR specific to influenza.

**Figure S6:**
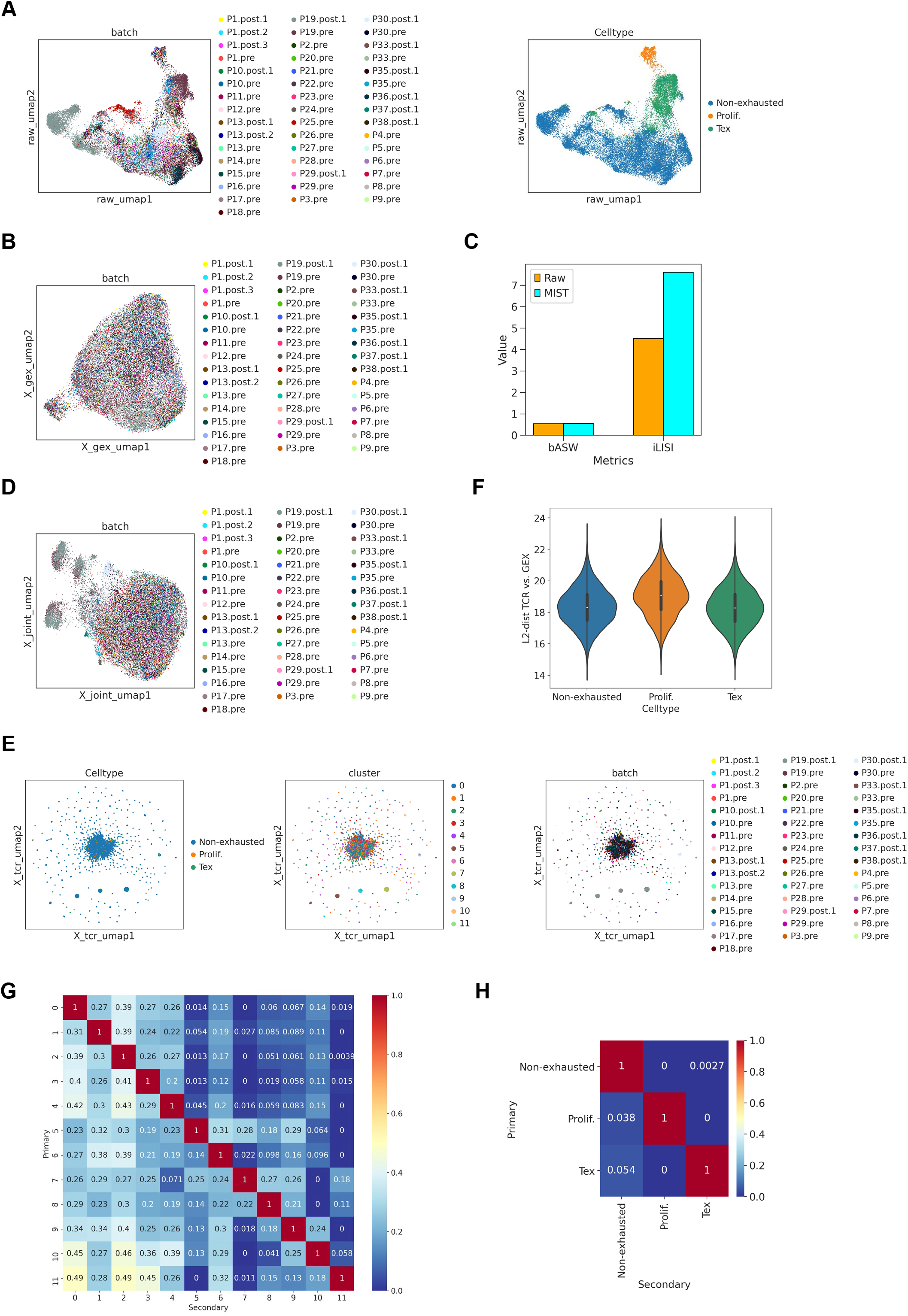
Analysis of T cells from NSCLC patients with anti-PD-1 therapy by the MIST model. A. UMAP plots of T cells in the anti-PD-1 therapy dataset by raw expression, color-coded by sample ID (left) and cell phenotype (right), respectively. Sample ID is designated as patient ID followed by anti-PD-1 therapy condition, with the numerical suffix indicating different sample replicate. Pre: before therapy; Post: after therapy. B. UMAP plots of T cells in the anti-PD-1 therapy dataset by the GEX latent representation from MIST, color-coded by sample ID. C. Statistical comparison of cell cluster robustness for the NSCLC anti-PD-1 therapy dataset when raw expression or GEX latent space from MIST were used for clustering. bASW: batch Average Silhouette Width; iLISI: integration Local Inverse Simpson’s Index. D. UMAP plots of T cells in the anti-PD-1 therapy dataset by the joint latent representation from MIST, color-coded by cell cluster ID. E. UMAP plots of T cells in the anti-PD-1 therapy dataset by the TCR latent representation from MIST, color-coded by cell phenotype (left), cluster ID (mid) and sample ID (right) and, respectively. F. Violin plots showing the distribution of Euclidean distances between the GEX and TCR latent representations for each T cells with distinct cell phenotype in the anti-PD-1 therapy dataset. G. TCR similarity between T cell clusters based on inter-cell Euclidean distance. H. Same as G, but for T cell phenotype.

**Figure S7:**
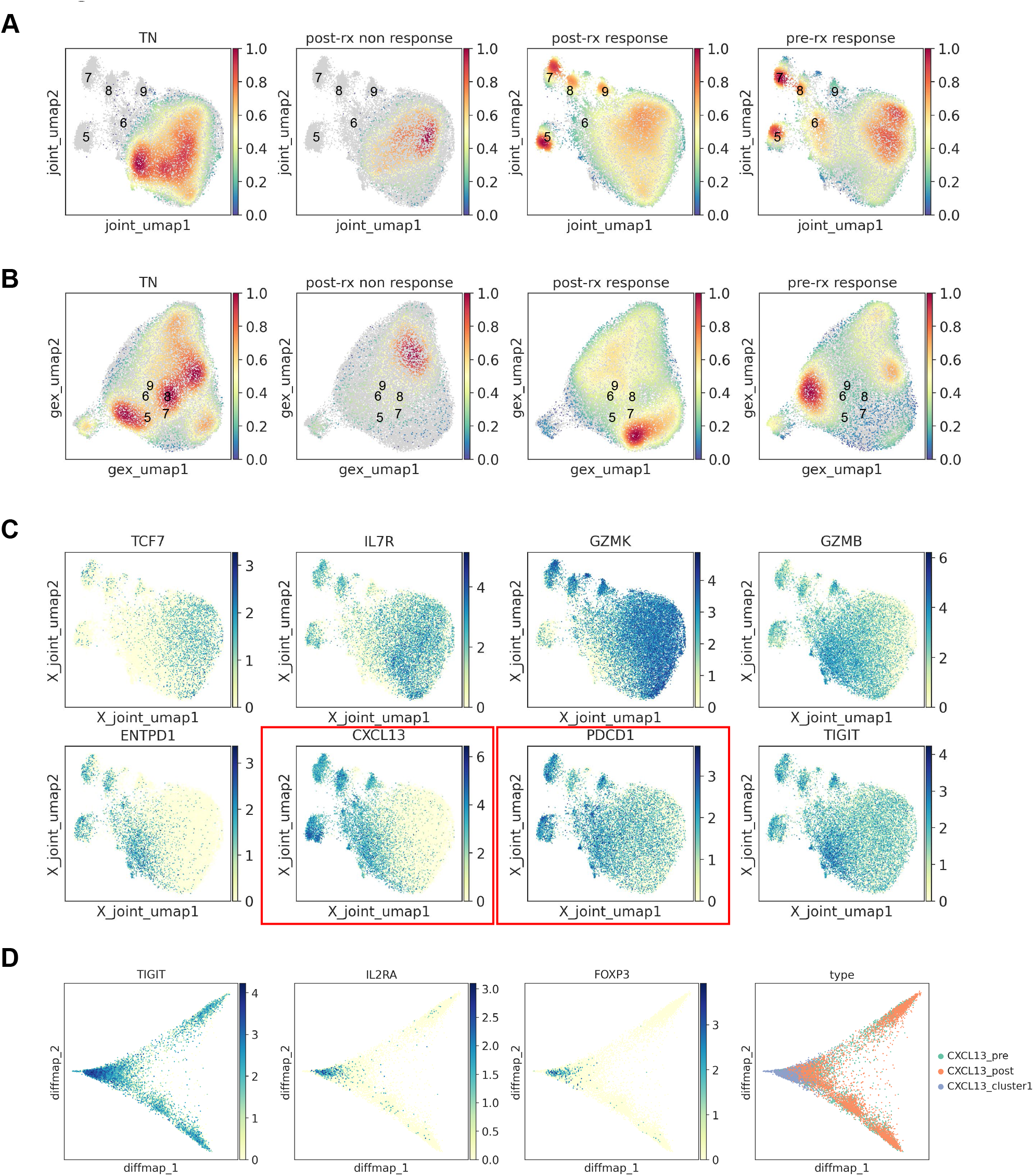
Additional characterization of T cells in the anti-PD-1 therapy dataset. A. UMAP plots of T cells in the anti-PD-1 therapy dataset by the joint latent representation from MIST, with color reflecting density of cells from different sample categories. TN: treatment naive; post-rx non-response: post-treatment with no response; post-rx response: post-treatment with response; pre-rx response: pre-treatment with response. B. UMAP plots of T cells in the anti-PD-1 therapy dataset by the GEX latent representation from MIST, with color reflecting density of cells from different sample categories. C. UMAP plots of T cells in the anti-PD-1 therapy dataset by the joint latent representation from MIST, with color reflecting expression level of the gene on top of each plot. D. From left to right: diffusion map visualization of CXCL13^+^ T cells, color-coded by raw expression level of TIGIT, IL2RA and FOXP3, and cell type, respectively.

## Notes

https://github.com/aapupu/MIST

### Competing Interest Statement

The authors have declared no competing interest.

### Summary of Updates

Updated reference citation and page number.

